# Abundance and community composition of comammox bacteria in different ecosystems by a universal primer set

**DOI:** 10.1101/492488

**Authors:** Zhirong Zhao, Guohe Huang, Mingyuan Wang, Nan Zhou, Shishi He, Chenyuan Dang, Jiawen Wang, Maosheng Zheng

## Abstract

Complete ammonia oxidizing bacteria (CAOB) have been recognized as a new member of ammonia-oxidizing microorganisms (AOMs) due to its single-step nitrification capability. However, the abundance and diversity of CAOB in environmental ecosystems were still far from known owing to the lack of specific molecular marker. Herein, a universal primer set specifically targeting the both clades of CAOB *amoA* gene with high specificity and coverage was successfully designed. Intriguingly, real-time quantitative PCR tests revealed that CAOB were ubiquitous and unexpectedly abundant in agricultural soils, river sediments, intertidal zones, drinking water and wastewater treatment systems. Phylogenetic analysis indicated that clade A existed in all the five types of ecosystems, while clade B were only detected in soil and sediment samples. Four sub-clusters were further classified within clade A, in which *N. nitrosa* cluster dominated CAOB *amoA* in activated sludge samples while the new recognized soil cluster was the primary constitute in soils. Moreover, the niche specialization between different CAOB species and the environmental conditions were supposed to be the primary driven force to shape the diversity and community of CAOB. This study provided a strong evidence in support of the ubiquities and high abundances of CAOB in various environmental ecosystems and highlighted the significance of including CAOB as the new member of AOMs to re-evaluate the biogeochemical nitrogen cycle.

## 1. Introduction

Nitrogen is an essential element for nucleic acids and proteins synthesis in all living organisms and the microorganism-mediated nitrogen cycle plays a critical role in the existence and sustainability of life. Ammonia oxidation to nitrite catalyzed by ammonia-oxidizing archaea (AOA) and bacteria (AOB) was recognized as the first and rate-limiting step of nitrification, followed by nitrite oxidation to nitrate by nitrite-oxidizing bacteria (NOB). This division of labor of ammonia oxidizing microorganisms (AOMs) and NOB has been long-held accepted as the only nitrification theory in the biogeochemical nitrogen cycle for more than 120 years since the discovery of AOB. Nevertheless, it was theoretically feasible to oxidize ammonia to nitrate by one single microbe, as complete ammonia oxidation (comammox) was more energy efficient than the traditional two-step nitrification process (Costa et al., 2006). This theory had not been experimentally testified until three complete ammonia-oxidizing bacteria (CAOB) capable of independently performing complete nitrification were enriched (Daims et al., 2015; Kessel et al., 2015).

Intriguingly, based on 16S rRNA gene all the three CAOB strains were phylogenetically affiliated with genus *Nitrospira*, which was the most abundant and widespread genus of NOB (Daims et al., 2016; Koch et al., 2015). However, CAOB were confirmed to possess a whole set of nitrifying genes encoding for ammonia monooxygenase (AMO), hydroxylamine dehydrogenase (HAO) and nitrite oxidoreductase (NXR), providing strong evidences for the capability of performing complete ammonia oxidation to nitrate via nitrite (Daims et al. 2015; van Kessel et al. 2015; Camejo et al. 2017). CAOB *amoA* gene formed a distinct phylogenetic cluster with a low similarity to the canonical AOB *amoA*, suggesting it was an ideal phylogenetic biomarker to identify and quantify CAOB in complex microbial communities. In addition, CAOB *amoA* gene was divergent which could be divided into two monophyletic clades, clade A and clade B (Pjevac et al., 2017).

To date, widespread attentions had been attracted on comammox owing to their potential contribution in ammonia oxidation and significant role in nitrogen cycle. The wide distribution of CAOB have been suggested through metagenomic surveys in some environmental habitats, such as paddy fields, agricultural soils, freshwater habitats and wastewater treatment systems (Kessel et al., 2015). High proportions of CAOB *amoA* gene were found within all the copper-containing membrane-bound monooxygenase (CuMMO) in nine environmental samples through a high degenerate primer (Wang et al., 2017a). Due to the divergence of the new *amoA* gene, two sister PCR primer sets targeting respective clade A and B were designed and used to explore the abundance and diversity of CAOB in different ecosystems (Pjevac et al., 2017). A newly designed primer set specifically targeting the clade A of CAOB *amoA* gene revealed high abundances of CAOB in full-scale WWTPs (Wang et al., 2018). Likewise, a clade A targeted primer set was designed to quantify the CAOB abundances in various environments (Xia et al., 2018).

However, the numerical abundances and community composition of CAOB in different ecosystems were still far from known. To fill this gap, the universal PCR primer set targeting both clade A and clade B of CAOB *amoA* was successfully designed in this study. With the specific amplification, the abundances and diversity of CAOB in five types of environmental habitats were investigated through quantitative PCR and high-throughput sequencing technology, respectively. This study provided an efficient molecular tool, which would facilitate the follow-up researches on comammox.

## 2 Materials and methods

### 2.1 Primer design and evaluation

PCR primer pairs targeting both clade A and clade B of CAOB *amoA* gene were designed to detect the presence of CAOB in diverse environmental samples. Briefly, putative CAOB *amoA* sequences affiliated with clade A or clade B were downloaded from National Center for Biotechnology Information (NCBI) as the query sequences. After clustering at 0.99 similarity level, a database including 120 putative CAOB *amoA* sequences were used to design the primer set.

To evaluate the specificity of the newly designed primer, the *amoA* database containing 43876 sequences and *pmoA* database containing 44402 sequences were downloaded from FunGene database. Then the phylogenetic tree was constructed using R, MAFFT, Xshell and FigTree software. The sequences that did not classified into the two genes were deleted to construct the accurate *amoA* and *pmoA* database containing 30963 and 31044 sequences, respectively. Primer coverage under 0, 1, 2 mismatch were calculated using the program primersearch of EMBOSS v6.5.7.0.

### 2.2 Collection of environmental samples

Eighteen samples were collected from five types of environmental ecosystems including agricultural soil, river sediment, drinking water, intertidal zone and activated sludge samples. All samples were immediately transported to the laboratory in ice and stored at −20◻. Detailed descriptions and detailed parameters of these samples were listed in Table 1.

**Table 1.** The characteristics of the environmental samples used in this study.

| Sample types | Sample ID | Longitude and latitude | Sampling Date | pH | NH<sub>4</sub>-N<sup>a</sup> |
| --- | --- | --- | --- | --- | --- |
| Activated sludge | XJH | 40°01'25", 116°18'20" | Sep-16 | 7.25 | 29.8 |
|  | YQ | 40°27'18", 115°58'24" | Sep-16 | 7.76 | 21.3 |
|  | YC | 34°25'34", 112°26'10" | Oct-16 | 7.2 | 24.4 |
|  | YS | 34°43'10", 112°47'07" | Oct-16 | 8.0 | 50.0 |
|  | XT | 27°49'57", 112°54'09" | Oct-16 | 7.39 | 10.4 |
|  | BB | 32°55'54", 117°24'23" | Oct-16 | 7.5 | 27.1 |
| Agricultural soil | SD | 32°59'34", 114°01'46" | Oct-16 | 5.5 | 18.2 |
|  | JZ | 38°02'37", 115°02'14" | Oct-16 | 6.5 | 10.0 |
|  | GZ | 37°04'36", 115°07'56" | Oct-16 | 5.5 | 7.7 |
|  | SH1 | 31°16'10", 121°26'29" | Oct-16 | 5 | 15.8 |
| River sediment | SZ | 22°33'44", 114°03'17" | Aug-17 | N.A. | N.A. |
|  | LZ | 36°04'47", 103°49'32" | Oct-16 | 7.54 | 5.4 |
|  | SMX | 34°47'50", 111°13'04" | Oct-16 | 7.38 | 3.8 |
| Drinking water | JN | 35°26'05", 116°35'05" | Oct-17 | N.A. | N.A. |
|  | BJ | 40°07'00", 116°14'01" | Sep-17 | N.A. | N.A. |
| Intertidal zone | SH | 31°20'24", 121°39'27" | Jul-17 | N.A. | N.A. |
|  | SZ1 | 22°30'47", 114°02'42" | Jul-17 | N.A. | N.A. |
|  | TJ | 39°06'04", 117°44'26" | Aug-17 | N.A. | N.A. |

<sup>a</sup> represents the influent ammonia concentration with unit of mg/L in activated sludge samples and mg/kg in agricultural soil samples.

### 2.3 DNA extraction and quantitative PCR

Total genomic DNA was extracted from each sample using the FastDNA Spin Kit for Soil (MP Biomedicals, USA) according to the manufacturer’s instructions.

The abundances of AOA, AOB and CAOB *amoA* genes were quantified with archamoA 19f/616r (Pester et al., 2012), *amoA* 1F/2R (Rotthauwe et al., 1997) and the present designed comamoA F/R on Applied Biosystems 7500 system, respectively. Quantitative PCR were performed in a 20 μL system consisted of 10 μL 2×SuperReal Premix Plus, 2 μL 50×Roxpreference Dye, 0.5 μL forward and reverse primers, 2 μL template DNA and 5 μL RNase-free H_2_O.

The AOA and CAOB *amoA* gene were amplified under the same condition: an initial denaturation step at 95 °C for 15 min, followed by 40 cycles of denaturation at 95 °C for 30 s, annealing at 53 °C for 30 s and extension at 72 °C for 1 min. For AOB *amoA* gene and 16S rRNA, the same condition except for annealing at 58 °C for 30 s and extension at 72 °C for 45 s was used. Each sample was conducted in three parallels. Standard curves were generated by ten-fold serially diluted plasmids containing the corresponding gene fragments as the template. Negative controls were conducted in which template DNA was replaced by RNase-free water. Melt curve and 1.5% agarose gel electrophoresis were used to confirm the specificity of the PCR products.

### 2.4 High-throughput sequencing and phylogenetic analysis

PCR amplicons of CAOB *amoA* were conducted with Illumina Miseq high-throughput Sequencing. The raw data were filtered and the remained sequences were clustered into operational taxonomic unit (OTU) at 97% nucleotide similarity with QIIME v1.9.1. The representative sequence of each OTU were aligned with the reference sequences collected from NCBI database to construct the Neighbor-Joining tree using 1,000 bootstrap replicates. Diversity indexes and rarefaction curves were also calculated at the OTU level. A principal coordinate analysis (PCoA) of weighted UniFrac distances between samples were calculated to address the community dissimilarities among the different environmental ecosystems. The raw high-throughput Sequencing data have been deposited at NCBI Short Read Archive (SRA) under BioProject PRJNA509063 and BioSample accession numbers SAMN10536240-10536256.

### 2.5 Statistical analysis

Analysis of variance (ANOVA) with Tukey test were carried out to compare different gene abundances and P < 0.05 was considered statistically significant.

## 3 Results

### 3.1 The specificity of the primer comamoA F/R

After in-silico and experimental confirmation, the forward primer comamoA F (5’-AGGNGAYTGGGAYTTCTGG-3’) and reverse primer comamoA R (5’-CGGACAWABRTGAABCCCAT-3’) with the highest specificity and coverage were selected. The primer did not nonspecifically hit any of the 30963 *amoA* sequences and 31044 *pmoA* sequences at mismatch of 0 and 1 bases. Even at the mismatch level of 2 bases, the primer could only target 18 *amoA* and 4 *pmoA* sequences, demonstrating the great potential to precisely identify the new microbe in complex environments.

### 3.2 Abundances of CAOB *amoA* gene in the five ecosystems

The abundances of CAOB *amoA* gene in the five types of environmental samples were determined via the newly designed primer and the canonical bacterial and archaeal *amoA* genes were also quantified for comparisons (Fig. 1). Unexpectedly, ubiquitous and abundant CAOB *amoA* gene were found to be distributed in activated sludge, intertidal zone, river sediment, agricultural soil, and drinking water systems.

**Fig. 1.**
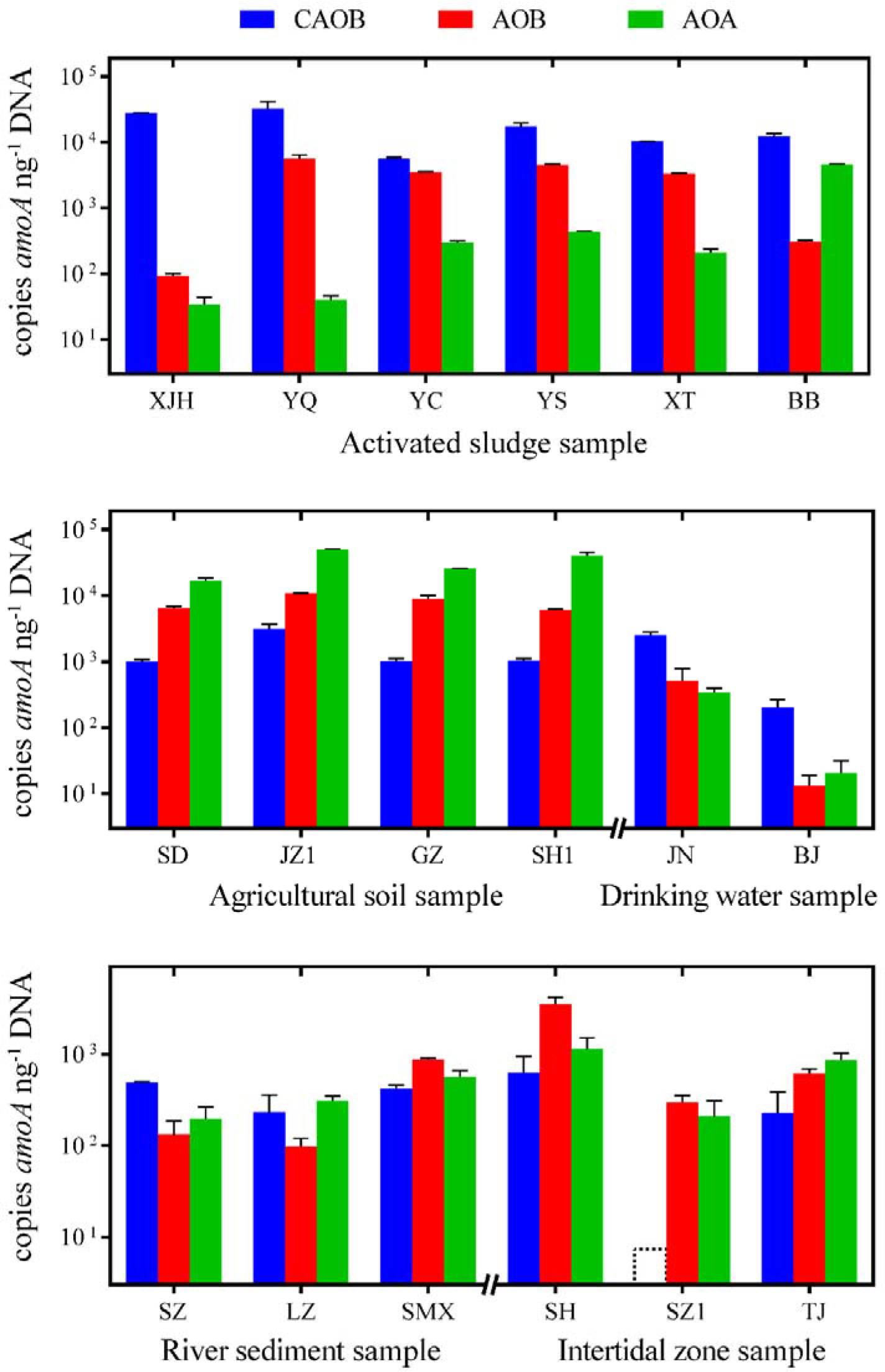
Abundances of *amoA* gene from CAOB, AOB and AOA in the 18 environmental samples. The dotted line in sample SZ1 represents below detection limit.

The abundance of CAOB *amoA* gene in activated sludge samples ranged from 5.5×10^3^ to 3.2×10^4^ copies ng^−1^ DNA, significantly higher than the canonical AOB *amoA* genes varying from 9.2×10^1^ to 5.6×10^3^ copies ng^−1^ DNA (P < 0.01, Fig. S1). Especially in sample XJH and BB, the ratio of CAOB to AOB *amoA* genes reached to 303.65 and 39.79, respectively. Comparing with the two bacterial *amoA* genes, the archaeal *amoA* gene always showed low abundances with an exception of sample BB. The numerical dominances of CAOB over AOB *amoA* also appeared in the two drinking water samples with 2.5×10^3^ and 8.3×10^1^ copies ng^−1^ DNA in JN and BJ, respectively. In contrast, opposite results were obtained in soil samples that AOA dominated among the three AOMs with 1.7×10^4^ to 4.8×10^4^ copies ng^−1^ DNA, while CAOB *amoA* gene were significantly lower with ratios of 0.03 ~ 0.60 to AOA *amoA* genes (P < 0.01). In the intertidal zone samples, the abundance of CAOB *amoA* gene in SZ was below detection limit and in SH and TJ they were as low as 3.2×10^2^ and 1.6×10^2^ copies ng^−1^ DNA, respectively. In river sediment, the abundances of CAOB *amoA* genes also ranged at relatively lower level and showed no significant differences with the canonical bacterial and archaeal *amoA* genes.

### 3.3 Diversity and phylogeny of CAOB *amoA* gene

To explore the community structure, high-throughput sequencing was conducted through the newly developed universal PCR primer set targeting both clade A and clade B. A total of 161,723 sequences were retrieved from the 17 environmental samples and 78 OTUs were clustered excluding the intertidal zone sample SZ1 which was failed in the amplification. The rarefaction curve and Good’s coverage (> 0.999) indicated the sequencing was nearly to saturation in all the environmental samples (Fig. S2). A neighbor-joining tree comprising 36 OTU representative sequences with proportion higher than 0.5% was constructed (Fig. 2).

**Fig. 2.**
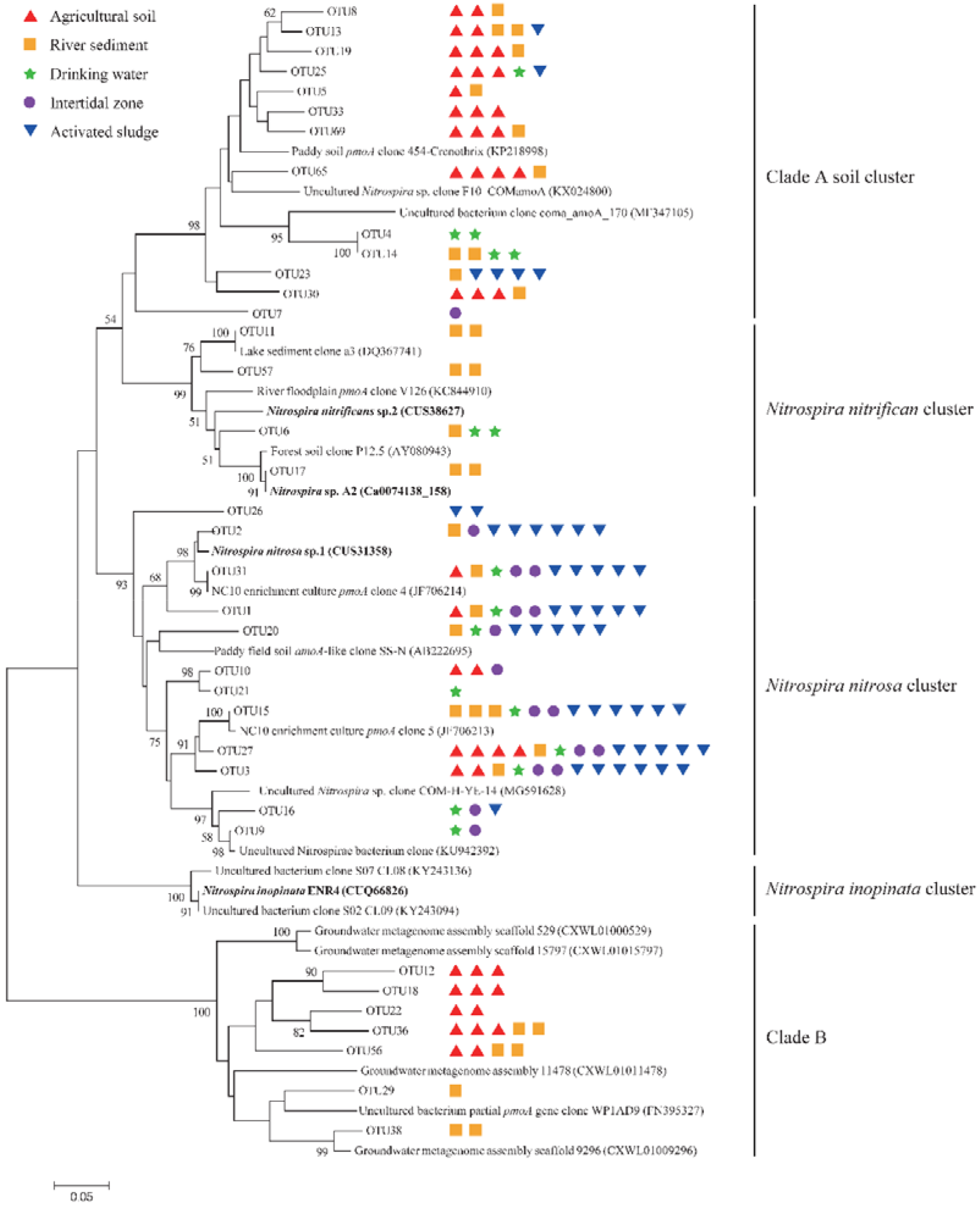
Neighbor-joining consensus tree generated from an alignment of CAOB *amoA* reference sequences and representative sequences of the top 36 OTU with proportion higher than 0.5% retrieved from the environmental samples. The bootstrap values higher than 50% are indicated at branch points. The scale bar represents 5% nucleic acid sequence divergence.

The obtained sequences formed a distinct cluster from the canonical bacterial and archaeal *amoA*, indicated the specificity of amplification by the current primer comamoA F/R. Consistent with previous results, the sequences were branched into two clades, where 64 out of the 78 OTUs were affiliated with clade A and the other 14 OTUs fell into clade B. The clustering analysis with heatmap visualization revealed the OTU distribution varied significantly between different types of environmental samples (Fig. 3). Moreover, four monophyletic sub-clusters were proposed to be further classified within clade A, among which three clusters were named as *N. nitrificans*, *N. inopinata*, *N. nitrosa* after the first cultured strain names and another one were tentatively named as clade A “soil” cluster given that majority of sequences in this cluster were retrieved from soil environmental samples. Moreover, the phylogenetic tree clearly showed that only clade A was encompassed in activated sludge, intertidal zone and drinking water samples and clade B was only identified in soil and sediment samples (Fig. 4).

**Fig. 3.**
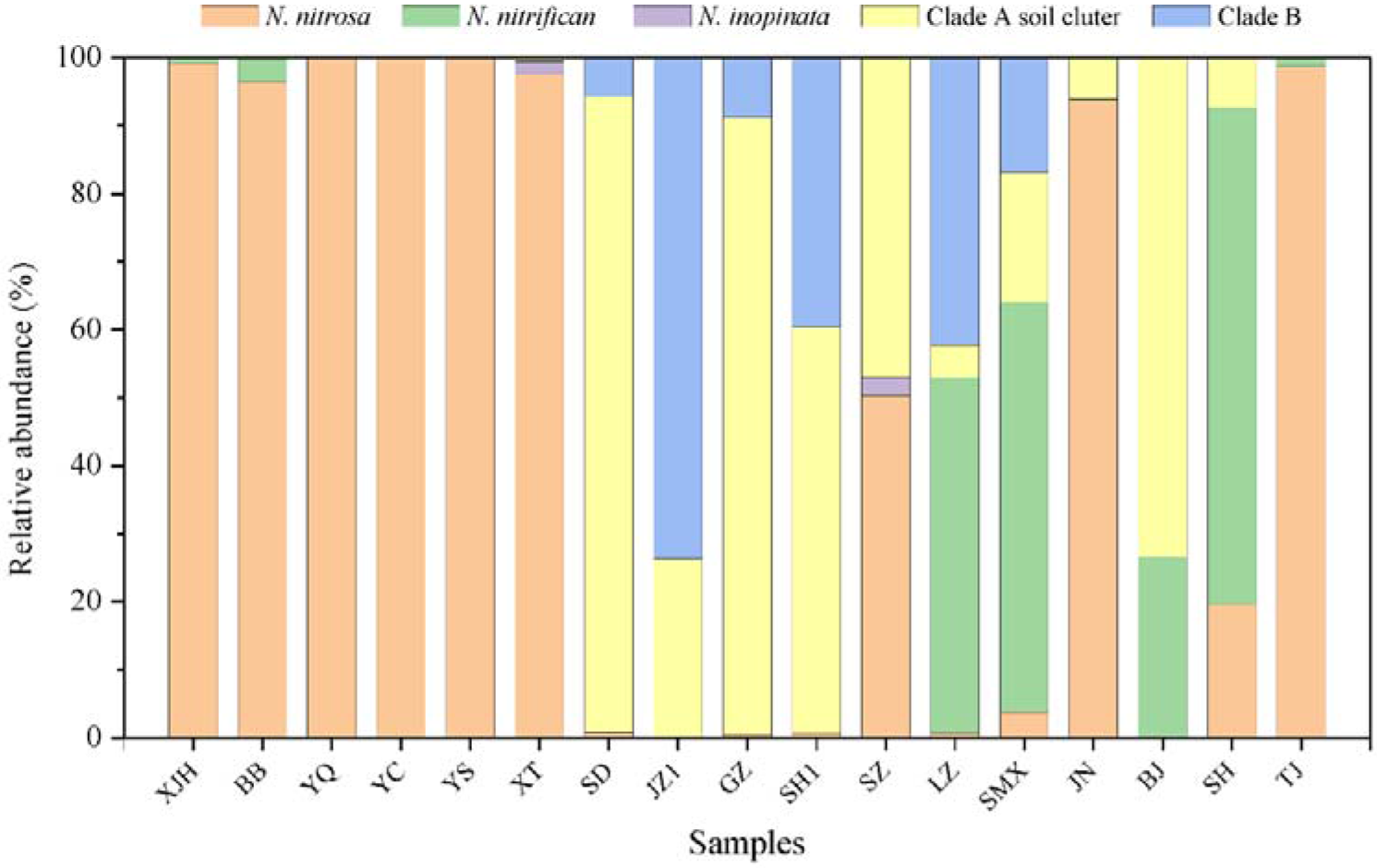
Distributions and relative abundances of different phylogenetic clusters of CAOB *amoA* in the 17 environmental samples. The amplification of intertidal zone sample SZ was failed and not shown.

**Fig. 4.**
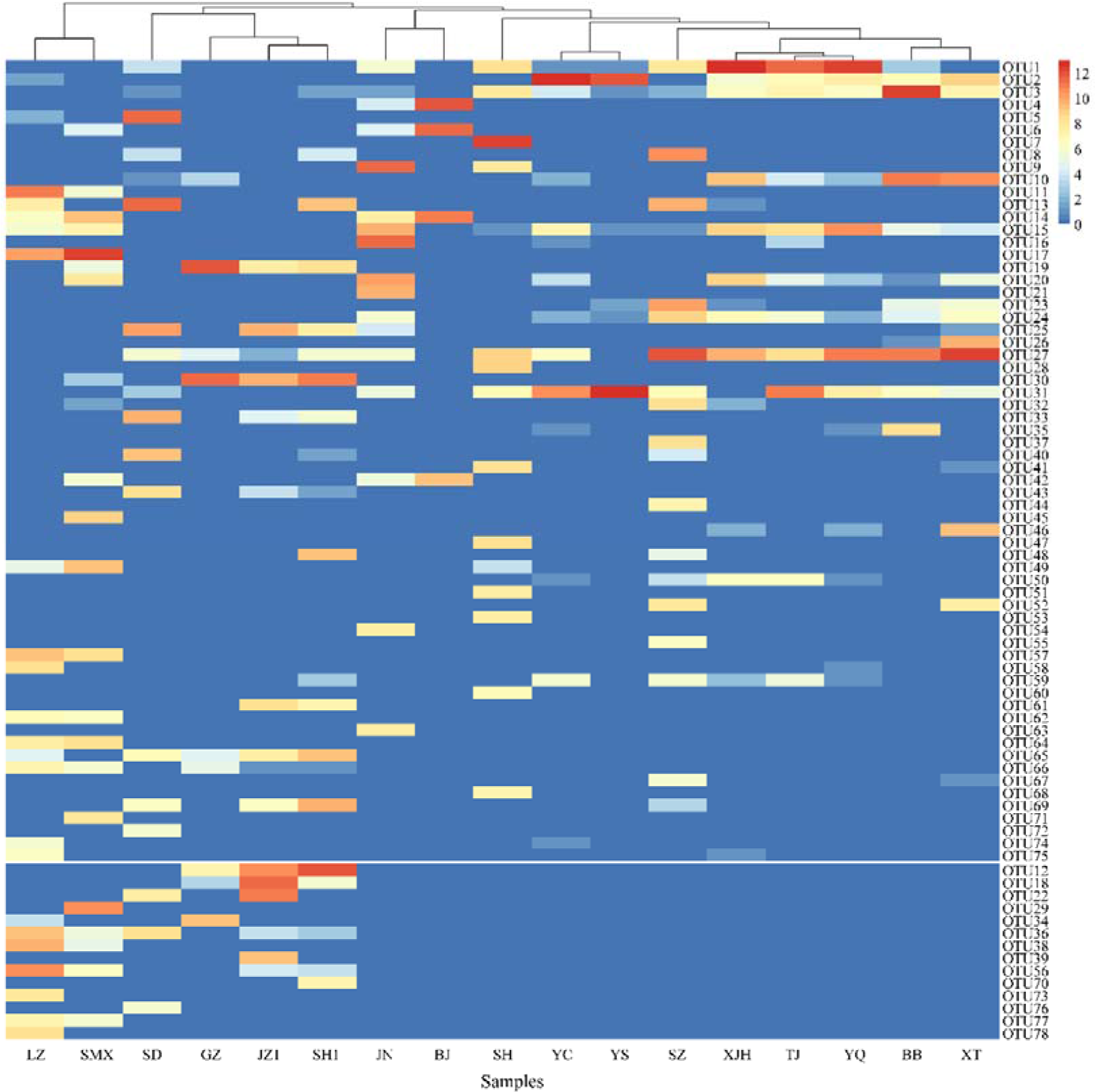
Heatmap displying the relative abundances of the 78 OTUs retrieved from the environmental samples. OTUs affilated with clade A and clade B were depicted above and below the white line respectively.

At the sub-cluster level, 14 OTUs retrieved from activated sludge were affiliated with cluster *N. nitrosa* which accounted for more than 98.8% of the total sequences, while the other three clusters only took small proportions in all the activated sludge samples, especially cluster *N. inopinata* was only detected in sample XT with a proportion of 1.8%. In drinking water samples, 93.6% of the sequences retrieved from sample JN were assigned into cluster *N. nitrosa* while another sample BJ was dominated by the soil cluster (73.3%) and *N. nitrificans* cluster (26.7%). The divergent result was also obtained in the two intertidal zone samples that *N. nitrosa* cluster took up to 98.7% of the total sequences in sample TJ whereas *N. Nitrificans* cluster (73.1%) and *N. nitrosa* cluster (19.75%) dominated in sample SH. The cluster *N. inopinata* was not existed in drinking water and intertidal zone samples.

In sharp contrast to activated sludge, all the soil samples harbored clade B sequences with proportion ranging from 5.7% to 73.6% and clade A was dominated by the soil cluster instead of cluster *N. nitrosa* and *N. nitrificans*. PCoA analysis revealed the significant clustering of the four soil samples from other environmental samples (Fig. 5). In the river sediment samples, LZ and SMX showed similar diversity in which clade B accounted for 42.2% and 16.9% of their respective sequences, whereas *N. nitrifican* cluster dominated within clade A. However, another sample SZ was distinct from LZ and SMX but it showed a much similar structure to activated sludge instead, which was probably attributed to the high ammonia concentration in the severely polluted river in which sample SZ was collected, demonstrating the importance of environmental factors in shaping the microbial community structure.

**Fig. 5.**
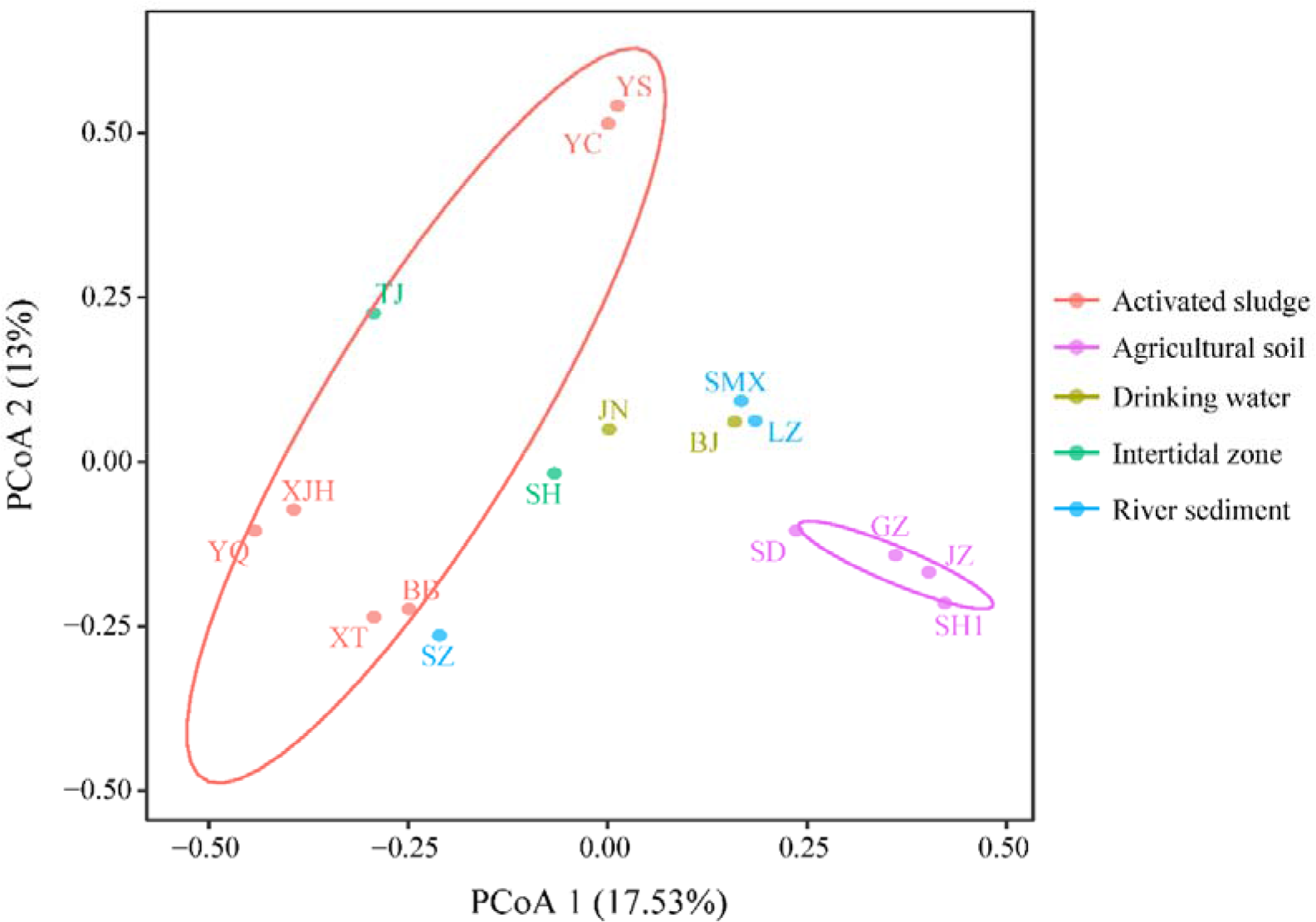
Principal coordinates analysis (PCoA) ordinations of the CAOB *amoA* community of the 17 environmental samples. Data points are colored by the different types of environmental ecosystems.

## 4 Discussion

### 4.1 Primer pairs targeting CAOB *amoA*

The discovery of comammox radically upended the century-old concept of the two-step nitrification process where ammonia-oxidizing bacteria and archaea mediated ammonia conversion to nitrite. Therefore, the identification and quantification of CAOB is of great significance to comprehensive and profound understanding of the nitrogen cycle. As CAOB do not form a monophyletic group within *Nitrospira* lineage II, they cannot be distinguished from the strict nitrite-oxidizing *Nitrospira* with 16S rRNA based analysis (Pjevac et al., 2017; Koch et al., 2018). Alternatively, the functional *amoA* gene formed a distinct group from the canonical AOB *amoA*, which become the best candidate of the specific molecular marker to identify CAOB in complex ecosystems.

Several primer sets have been developed for quantifying CAOB *amoA* gene using quantitative real-time PCR analyses since their discovery. The divergence of CAOB *amoA* hindered the development of the universal primer set covering both clades. The primer set targeting only clade A had been designed to quantify CAOB in wastewater treatment systems and other ecosystems (Wang et al., 2018; Xia et al., 2018). The sister primer sets target clade A and clade B were separately developed for identifying CAOB, although nonspecific amplifications were reported (Pjevacet al. 2017; Keene-Beach and Noguera, 2018). Therefore, it was urgent to develop a universal primer pair targeting both clade A and clade B with a higher degeneration to expand the application in various environmental habitats. Herein, the present new universal primer set comamoA F/R possessing high specificity and coverage of CAOB *amoA* was developed and confirmed by in silico analysis and high-throughput sequencing, which provided an effective and efficient tool for detecting and quantifying CAOB in complex ecosystems in the follow-up comammox related research.

It was noteworthy that the primer set A189/A682 previously designed for detecting *pmoA* of methane-oxidizing bacteria (MOB) could amplify a proportion of the new recognized *amoA* gene due to their close phylogenetic relationship (Dörr et al., 2010; Ho et al. 2016; Holmes et al., 1995). The nonspecific amplification resulted in that hundreds of sequences affiliated with CAOB *amoA* had been deposited in public NCBI database and incorrectly classified as canonical *pmoA* or *amoA* sequences before the discovery of comammox. Likewise, a number of CAOB *amoA* sequences were also included in the *amoA* and *pmoA* database from FunGene database, suggesting the necessity of filtering the CAOB *amoA* sequences to ensure the accuracy of the database. In addition, the primer set A189/A682 to identify MOB should be used with caution to avoid the nonspecific amplification of CAOB *amoA* in the future researches.

### 4.2 The ubiquity and diversity of CAOB *amoA* in various ecosystems

Ammonia oxidation was recognized as the central and rate-limiting step of nitrogen cycle in environmental ecosystems, which was considered to be driven by the two counterparts AOB and AOA before the discovery of CAOB. Their abundances and activity had been extensively studied to explore their respective role in ammonia oxidation in various ecosystems (Zheng et al., 2017). However, the discovery of CAOB raised new questions regarding their abundances and functions in driving ammonia oxidation, especially comparing with the canonical ammonia oxidizers. Intriguingly, the present results revealed that CAOB *amoA* genes had significantly higher or comparable abundances with AOB and AOA *amoA* genes in most environmental samples, indicating they were probably an overlooked but significant ammonia oxidizer actively participating in ammonia oxidation.

In WWTPs, AOB was always proposed to be the main contributor to ammonia oxidation due to their dominant numerical abundances over AOA. In the present study similar results were also obtained in most samples with an exception of sample BB that encompassed more abundant archaeal *amoA*. The preponderance of AOA could be ascribed to the niche specialization of AOA preferring low ammonia concentration, low pH value and antibiotic or petroleum-containing environment (Zhang et al., 2015; Mussmann et al., 2011; Pan et al., 2018). Unexpectedly, CAOB *amoA* genes presented higher abundances than AOB and AOA *amoA* genes in all the activated sludge samples, especially in sample XJH and BB the ratio of CAOB to AOB *amoA* genes reached up to 303.65 and 39.79, respectively. The prevalence of CAOB in activated sludge was also revealed through the metagenomic survey that CAOB-assigned coding regions accounted for 0.28 ~ 0.64% of total coding DNA sequences in 16 full-scale WWTPs (Annavajhala et al., 2018). The large proportion of CAOB in WWTP VetMed was supposed to be the primary contribution to the higher abundance of genus *Nitrospira* than canonical AOB (Daim et al., 2015). The considerable amount of CAOB among AOMs highlighted that they might take the primary responsibility for ammonia oxidation together with AOB (Zheng et al., 2018). The gradual oxygen and substrate concentration throughout the activated sludge flocs might be one important factor providing the suitable habitat favoring the CAOB propagation.

Drinking water systems was an ammonia-depleted ecosystem, the existence of ammonia in drinking water can be imported from the source water or artificial addition of chloramine disinfectants to produce biologically stable water for distribution (Regan et al., 2003; van der Wielen et al., 2009). The limited ammonia concentration in drinking water provided suitable substrate concentration for the attached biofilm growth although nitrification in the water distribution systems was unwanted to avoid the chloramine decay (Lawson and Lücker, 2018). An metagenomic analysis revealed the preponderance of CAOB than the canonical AOB and AOA in drinking water samples (Wang et al., 2017b). The high abundance of CAOB was also detected in rapid sand filtration (RSF) system for groundwater treatment (Palomo et al., 2016). It was tempting to speculate that the frequently found numerical dominance of *Nitrospira* over AOB and AOA in drinking water distribution systems was probably ascribed from the abundant CAOB benefiting from the attached growth on the sand surface provided by the special configuration of RSF system (Gülay et al., 2016; Tatari et al., 2017).

The intertidal zone ecosystem was the ocean-land interaction zone between the low-tide and high-tide line suffering the intense alternative exposure of air and seawater. The abundances of CAOB *amoA* in the intertidal samples were relatively low comparing with other ecosystems and became undetectable in the sample SZ. The intense alternative exposure to air would greatly suppress the propagation of CAOB ascribed to their niche preference of low-oxygen condition. Besides, ocean was proved to be not a suitable habitat as they were absent in oceanic samples, which was probably ascribed from the high salinity (Kuypers, 2017; Santos et al., 2017).

Nitrification process in agricultural soils would result in loss of nitrogen fertilizer and pollution of surrounding waters. The numerical dominances of AOA over AOB were found in majority of researches on soil ecosystems attributed to their higher ammonia affinity and adaption to acidic environments (Kits et al., 2017; Leininger et al., 2006; Nicol et al., 2008). Similar results were also obtained in the present study. CAOB were considered to possess similar ecological niche of low substrate concentration as they both had high affinity to ammonia. However, researches regarding the abundances and functions of CAOB in soil were still unclear. It was found that CAOB *amoA* sequences accounted for more than 20% of all the CuMMO encoded gene using a highly degenerate primer (Wang et al., 2017a). The high abundance of both CAOB *amoA* clade A and clade B were determined in 300 forest soil samples using the sister primer sets, suggesting they might functionally outcompete the canonical AOA under the oligotrophic soil ecosystems (Hu and He, 2017). However, AOA rather than CAOB were proved to conduct efficient autotrophic nitrification in the forest soil with ^13^C labelled CO_2_ microcosm experiment (Shi et al., 2018).

Notably, sequences affiliated with clade B were identified in all the soil samples with highest proportion up to 73.6%, in sharp contrast to activated sludge samples containing not any clade B assigned sequences. Instead, activated sludge samples were highly dominated by *N. nitrosa* cluster within clade A, which was consistent with a previous study on 19 WWTPs using a species-level targeted primer set (Keene-Beach and Noguera, 2018). Besides, the dominance of *N. nitrosa* cluster was also identified in the sediment of a heavily polluted river. Therefore, it seemed that *N. nitrosa* possess the ecological niche of high substrate concentration while clade B preferred to live in ammonia-depleted environments. The divergence might be attributed to the distinct membrane ammonium transporter families harbored by the two CAOB clades that clade A possess Rh-type transporters with low ammonia affinity and high uptake capacity while clade B possess Amt-type transporters with a relatively high ammonia affinity (Palomo et al. 2018). Moreover, the Rh-type ammonia transporters showed high homology (>70% amino acid similarity) to *Nitrosomonas europea* (van Kessel, et al., 2015). It was tempting to speculate that *N. nitrosa* cluster had the similar ecological niche with the canonical AOB species adapting to ammonia-rich environments whereas clade B preferred to oligotrophic environments.

The niche specialization and environmental parameters were the dominant factors in shaping the microbial community in various ecosystems (Bahram et al., 2018). However, it is still unclear to which extend the environmental factors such as the process configuration, nutrient concentration and biogeography influence the abundance and diversity of CAOB. Given the potential ecological significance of comammox in the nitrogen cycle, CAOB should be taken into account as one significant AOM in future studies to re-assess the niche specialization and relative contribution together with other nitrifying microorganism in various environments, which is crucial to get a panoramic view of the biogeochemical nitrogen cycle.

## 5 Conclusion

In this study, a primer set targeting both clade A and clade B of CAOB *amoA* gene was successfully designed. CAOB was found to be widely distributed with significantly higher or comparable abundances compared with canonical AOB and AOA in the investigated environmental samples. The monophyletic clade A and clade B were clustered in the CAOB *amoA* gene and four sub-clusters were further classified within clade A. The cluster *N. nitrosa* dominated CAOB *amoA* in activated sludge samples while clade B and the new recognized soil cluster within clade A were the primary constitutes in agricultural soils. The distinct communities were supposed to be shaped by niche specialization of different CAOB species and the environmental conditions in various ecosystems. This study provided an effective molecular tool for the following researches on comammox and offered new insights into the ubiquities and community compositions of CAOB in various environmental ecosystems.

## Supporting information

## Acknowledgements

This work was supported by National Natural Science Foundation [Grant No. 41701278] and Beijing Natural Science Foundation [Grant No. 8174075].

