## Supplementary material for "Abundance and community composition of comammox bacteria in different ecosystems by a universal primer set"

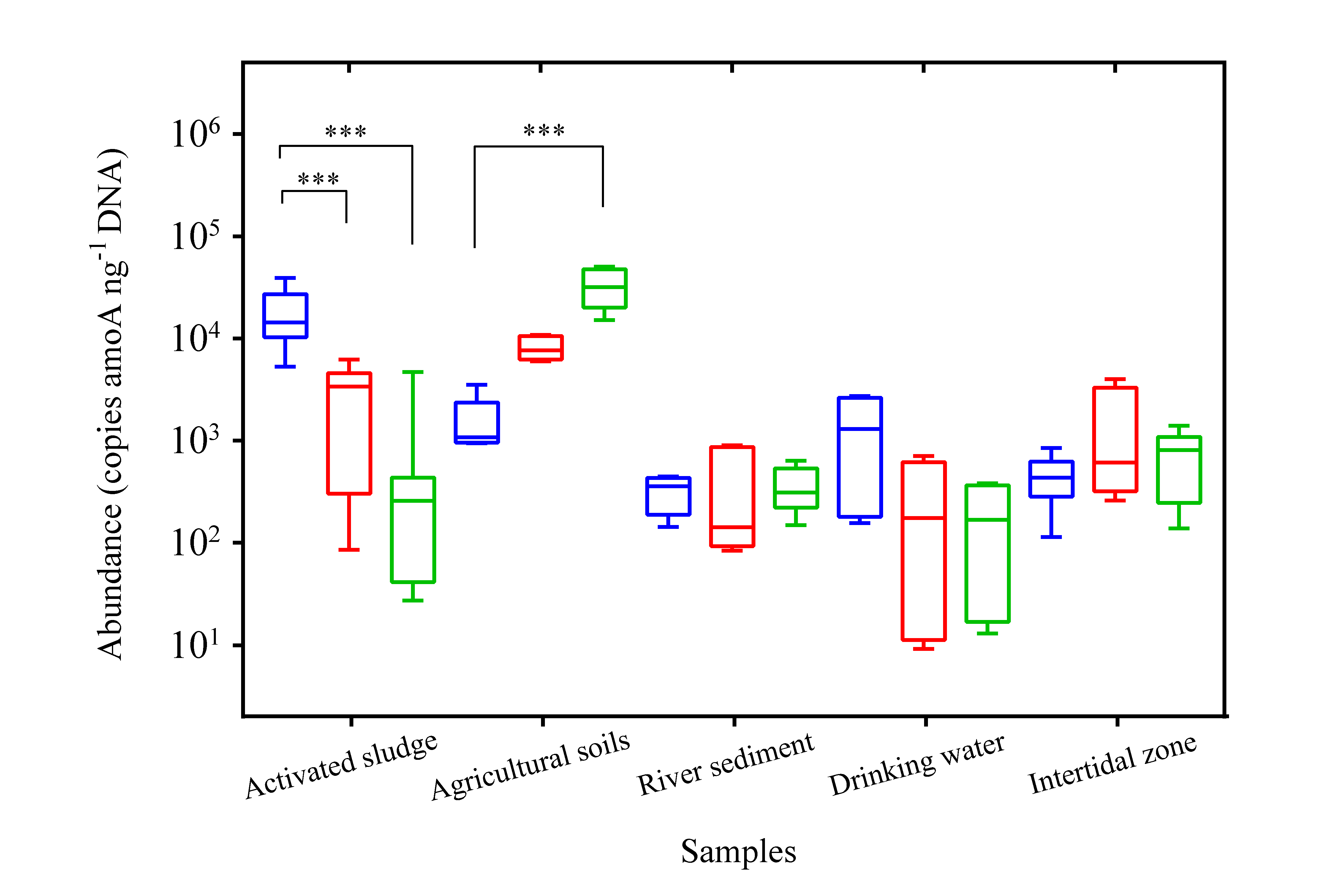

Fig. S1 Abundances of *amoA* gene from CAOB, AOB and AOA in the five environmental ecosystems. The blue, red and green box represents CAOB, AOB and AOA, respectively.

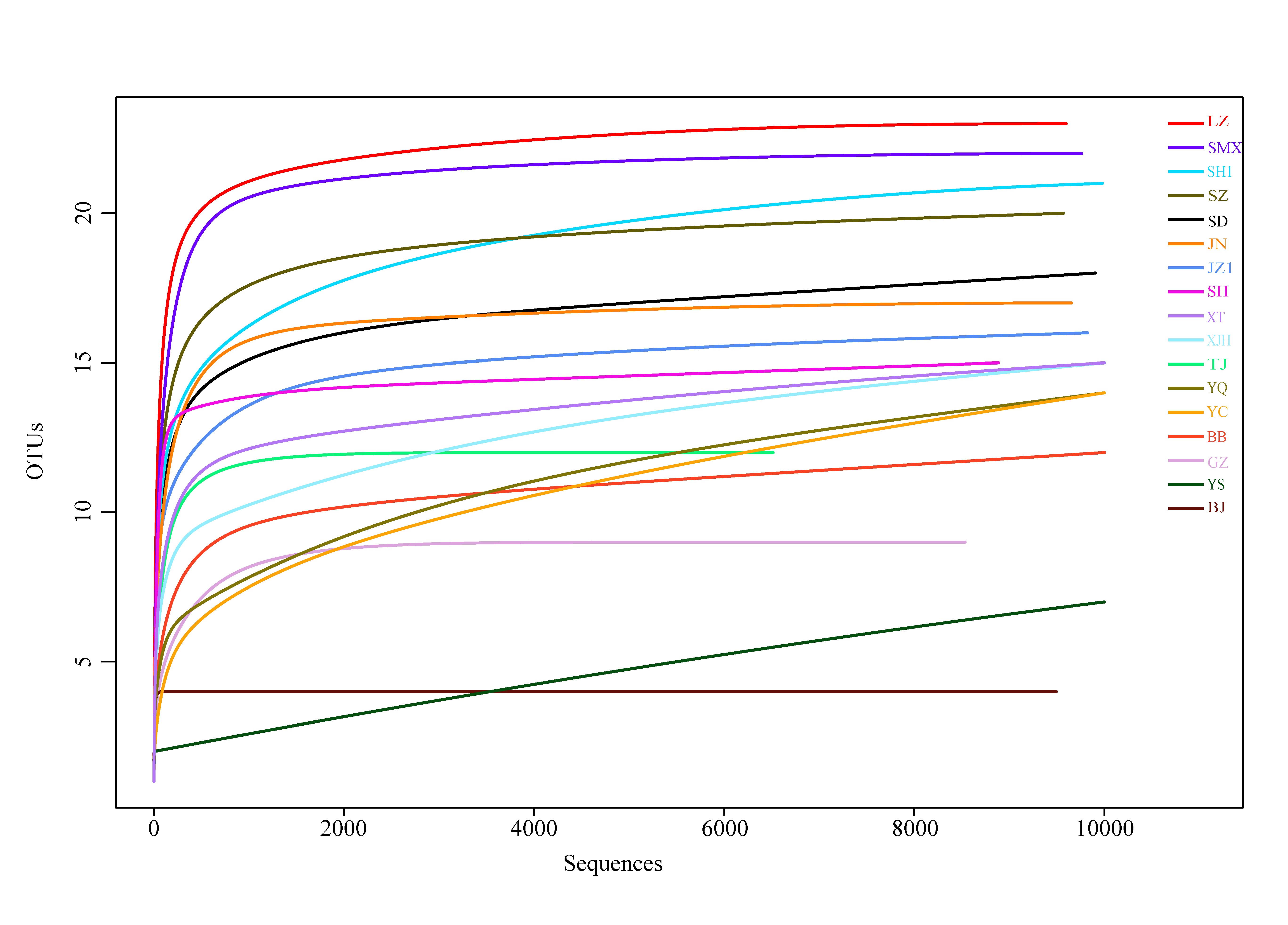

Fig. S2 Rarefaction curves of observed OTUs at 97% sequence similarities of the different samples.
